## Supplementary figures and tables for "Structural mechanism of strand exchange by the RAD51 filament"

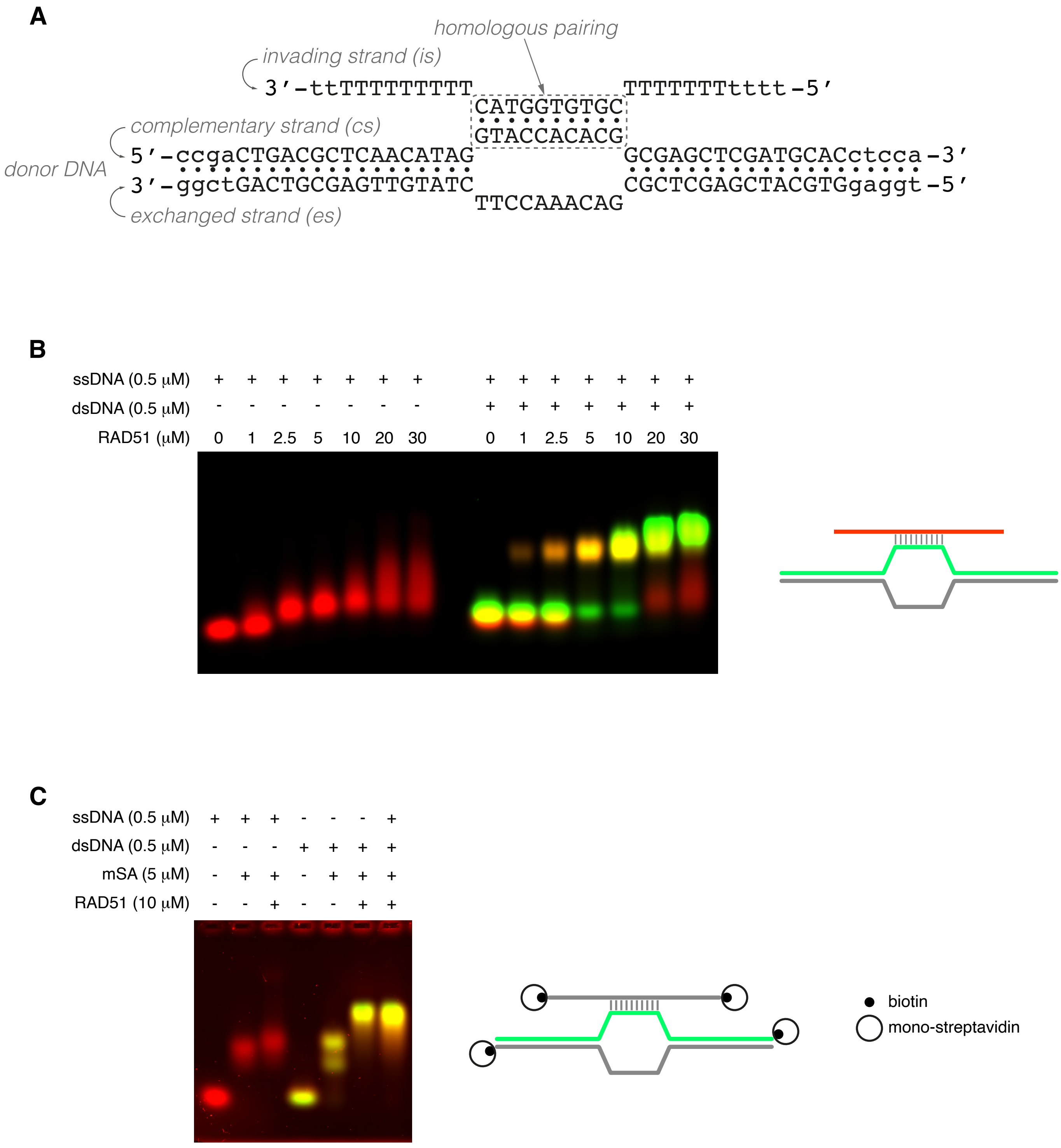


**Figure S1. D-loop reconstitution with human RAD51. A** Sequences and designed base pairing of oligonucleotides used in D-loop reconstitution for cryoEM analysis. Nucleotides in lower case were not included in the final atomic model. **B, C** Electrophoretic mobility shift assays (EMSAs), shown as composite fluorescent images: the Cy5-labelled (B) or SYBR Gold-stained ssDNA (B) is coloured red, while the Cy3-labelled strand in dsDNA is green. Schematic drawings of the DNA substrates coloured according to the fluorescence label are shown above each experiment. **B** RAD51 titration on ssDNA only or ssDNA and complementary dsDNA. **C** RAD51 binding to doubly-biotinylated ssDNA only or to doubly-biotinylated ssDNA and complementary doubly-biotinylated duplex DNA, in the presence of mono-streptavidin (mSA).

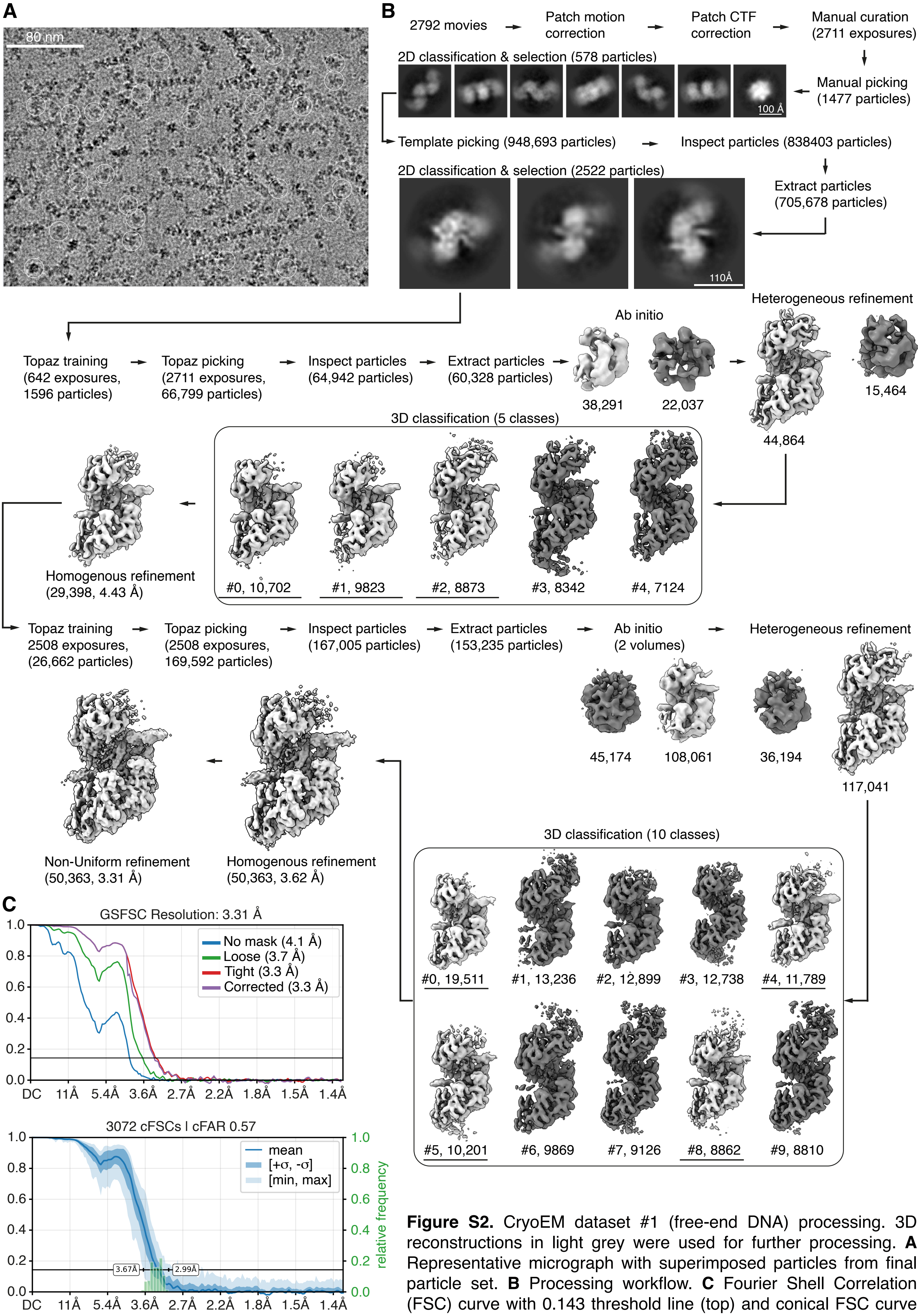


**Figure S2.** CryoEM dataset #1 (free-end DNA) processing. 3D reconstructions in light grey were used for further processing. **A** Representative micrograph with superimposed particles from final particle set. **B** Processing workflow. **C** Fourier Shell Correlation (FSC) curve with 0.143 threshold line (top) and conical FSC curve for directional resolution analysis (bottom).

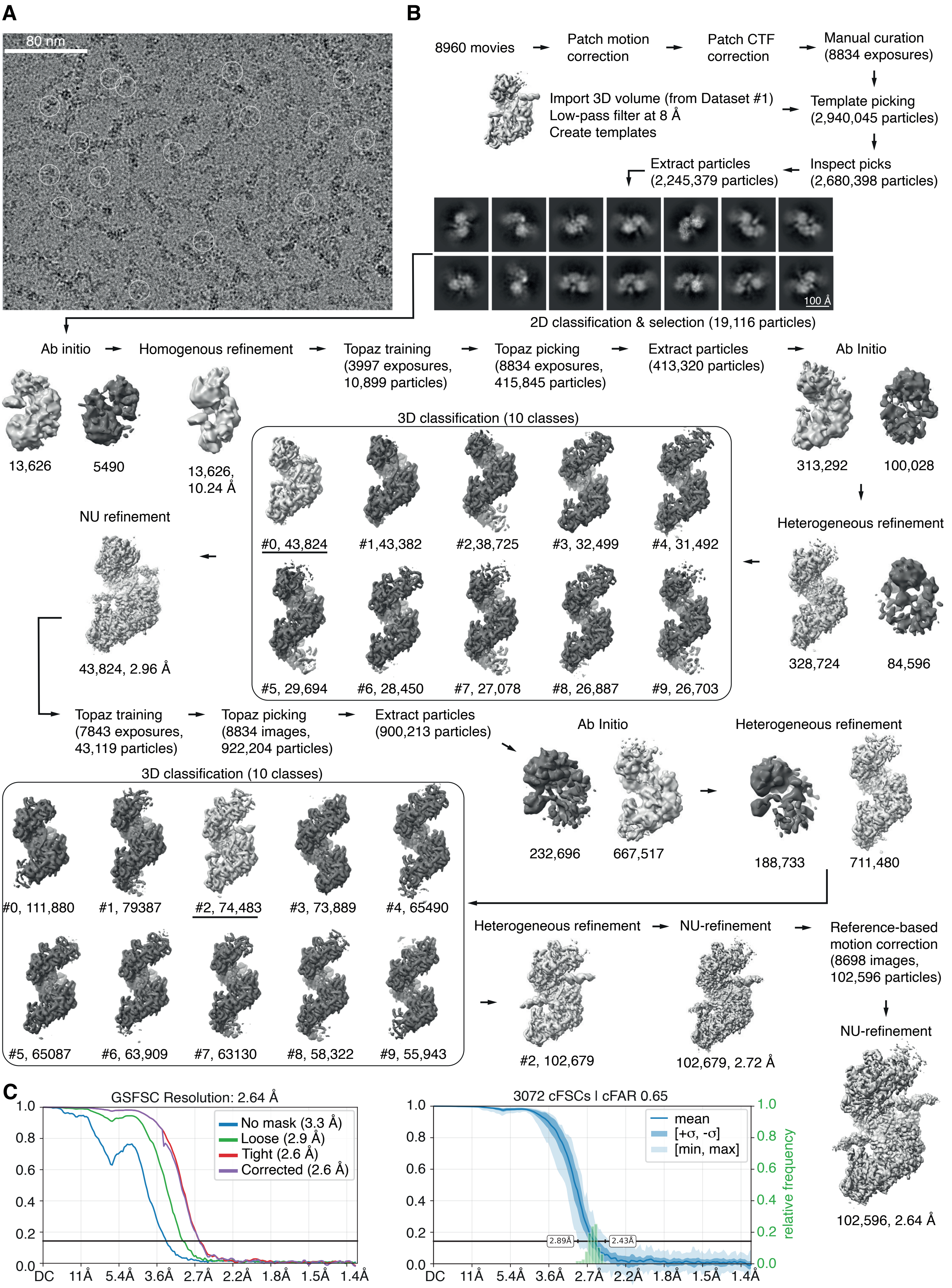


**Figure S3.** CryoEM dataset #2 (streptavidin-capped DNA) processing. 3D reconstructions in light grey were used for further processing. **A** Representative micrograph with superimposed particles from final particle set. **B** Processing workflow. **C** Fourier Shell Correlation (FSC) curve with 0.143 threshold line (left) and conical FSC curve for directional resolution analysis (right).

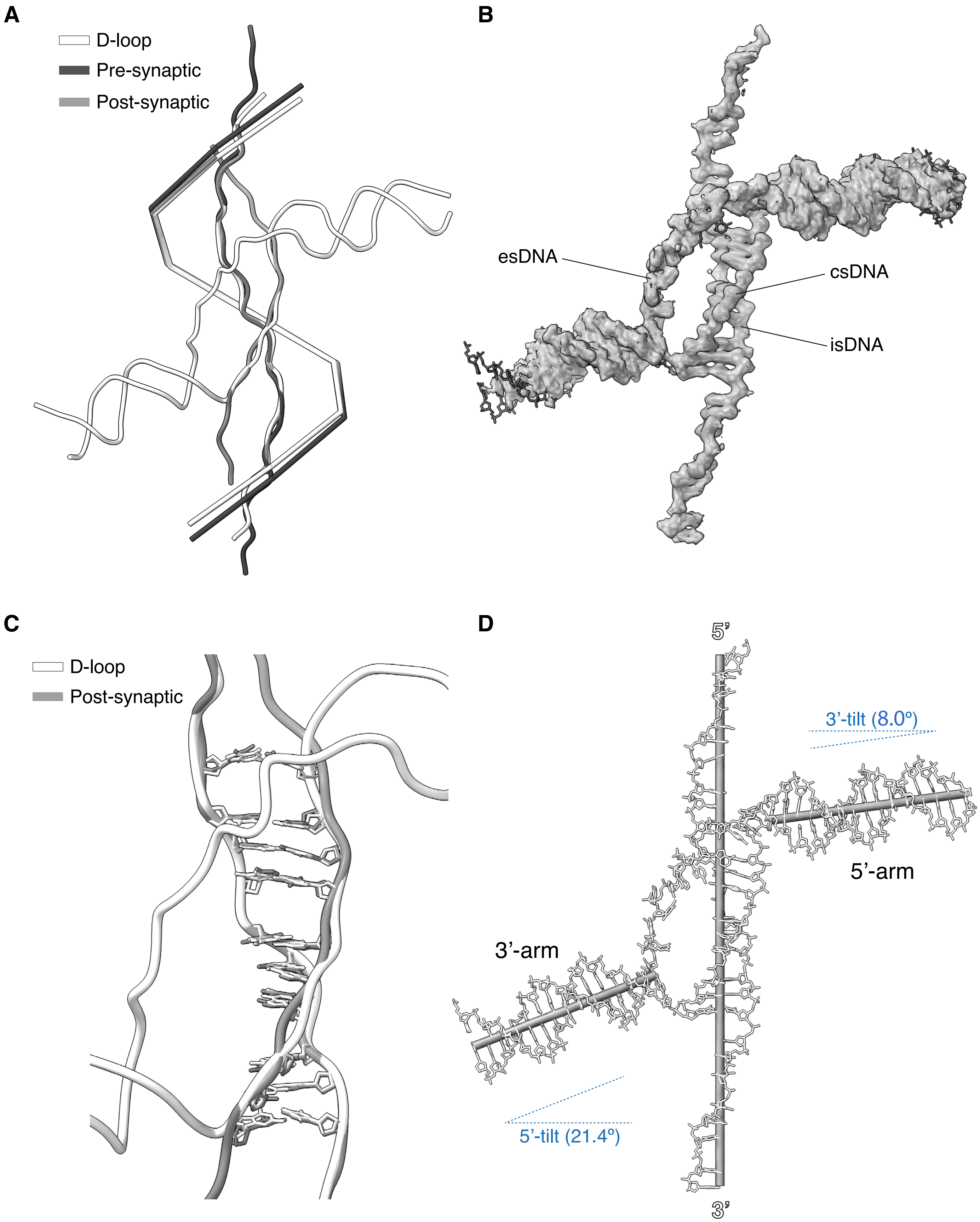


**Figure S4. Geometric analysis of D-loop DNA. A** Superposition of D-loop, pre-synaptic (PDB ID: 8BQ2) and post-synaptic (PDB ID: 8BR2) filament structures. Only the DNA is shown, as a tube ribbon coloured white (D-loop), grey (post-synaptic DNA) and dark grey (pre-synaptic DNA). The centroid positions of the RAD51 protomers in each structure are joined, to generate a trajectory representative of the filament pitch for each structure. **B** Details of the cryoEM map for the D-loop DNA, with superimposed atomic model of the DNA strands. **C** Superposition of D-loop and post-synaptic DNA, oriented and coloured as in panel A, showing the base pairs in the homologous pairing sequence of the D-loop and post-synaptic dsDNA. **D** Orientation of the axes of the arms of the donor dsDNA relative to the filament axis. The axes are drawn as cylinders and were fitted based on the centroid positions of the base pairs (for the donor arms) or the nucleotides of ssDNA in the filament. The tilt angles of the dsDNA arms relative to the filament axis are indicated.

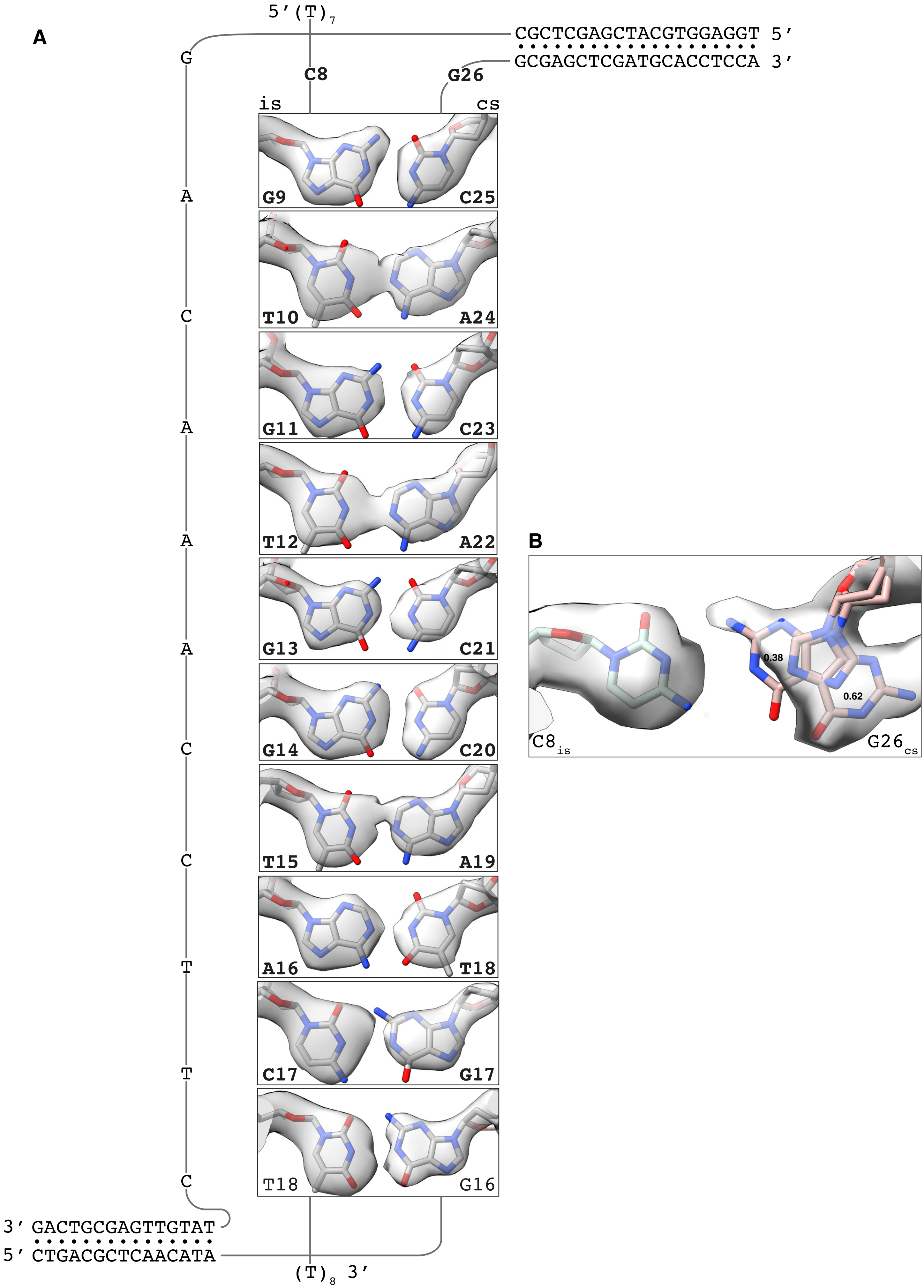


**Figure S5. CryoEM density for homologous base pairing.** Details of the cryoEM map for the base pairs formed by the invading strand (is) of the filament DNA with the complementary strand (cs) of the donor DNA. Nucleotides designed to base pair in the D-loop structure are in bold. **A** Homologous base pairing. **B** Base conformation for the designed $C8_{is}$ : $G26_{cs}$ pair. Occupancy values for the two G26 conformations are reported in the panel.

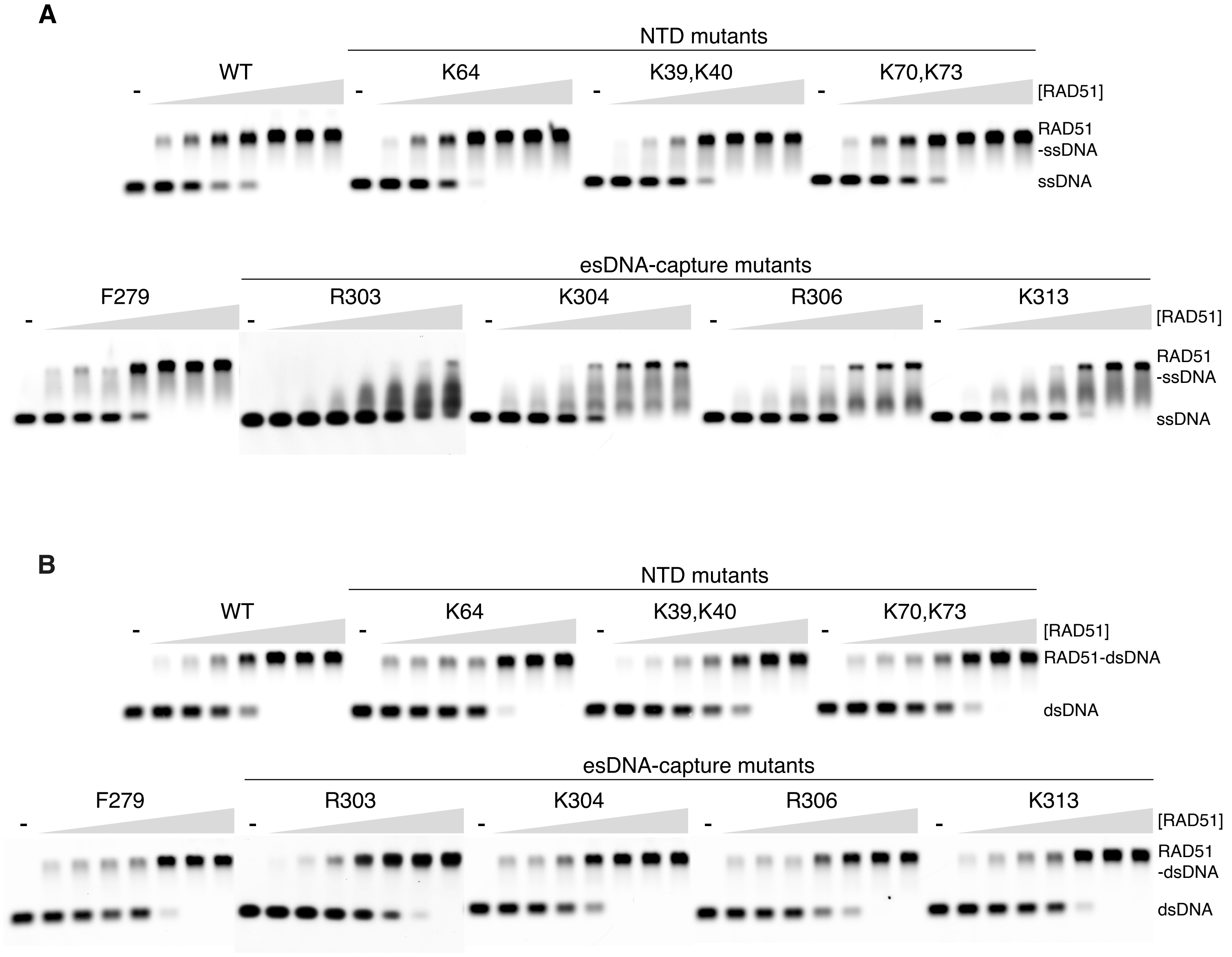


**Figure S6. Electrophoretic mobility shift assay for the interaction of RAD51 mutants with ssDNA (A) and dsDNA (B).** For each mutant, increasing proten concentrations (0.5, 1, 2, 4, 6, 8, 10 μM) were titrated against a fixed amount of DNA (0.5 μM).

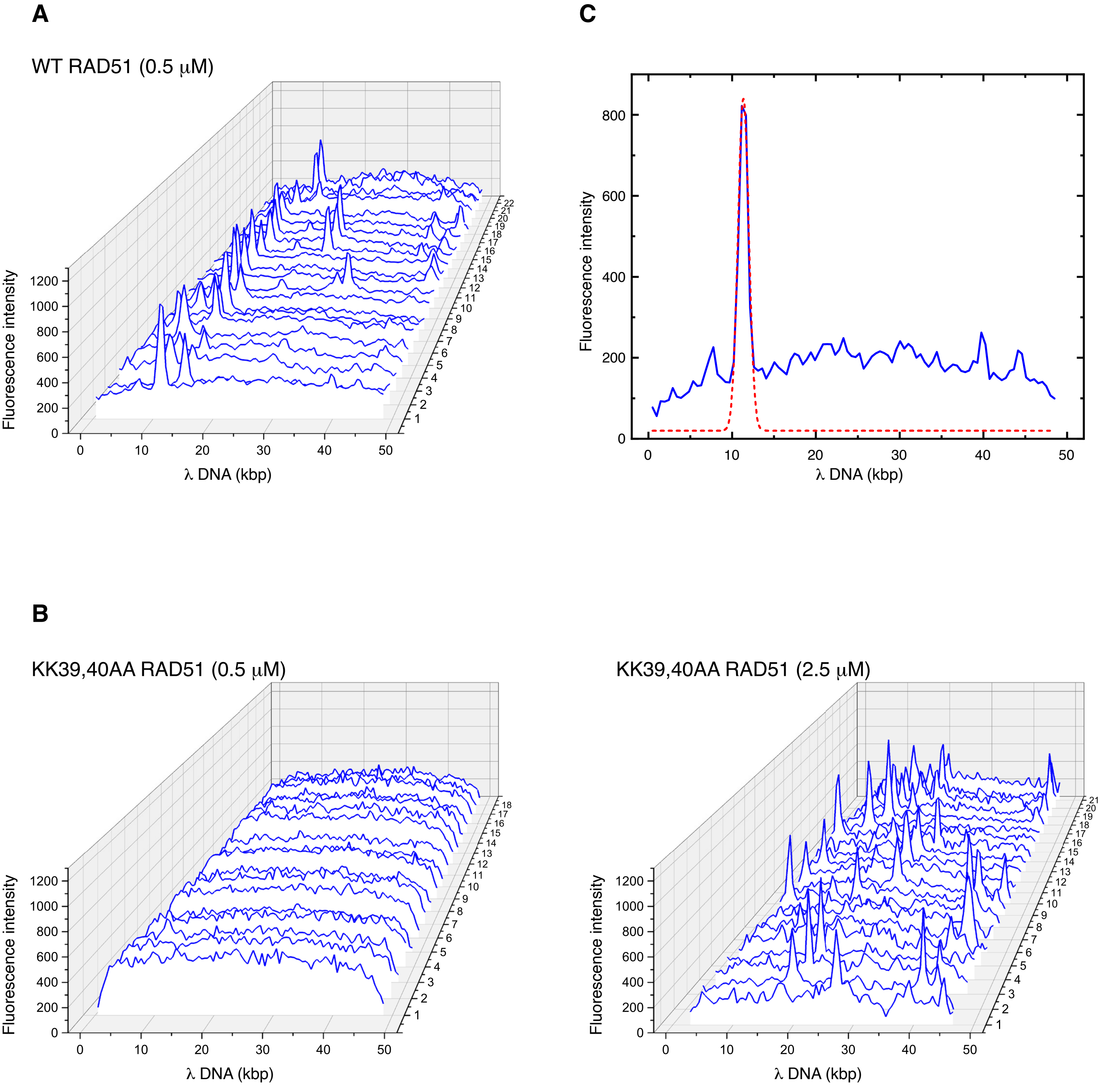


**Figure S7.** Fluorescence intensity traces extracted from 2D scans of trapped DNA molecules. **A** Traces for the wild-type RAD51 filaments. **B** Traces for KK39,40AA RAD51 mutant filaments, at 0.5 µM (left) and 2.5 µM (right).**C** Gaussian peak fitting for a representative trace of wild-type RAD51.

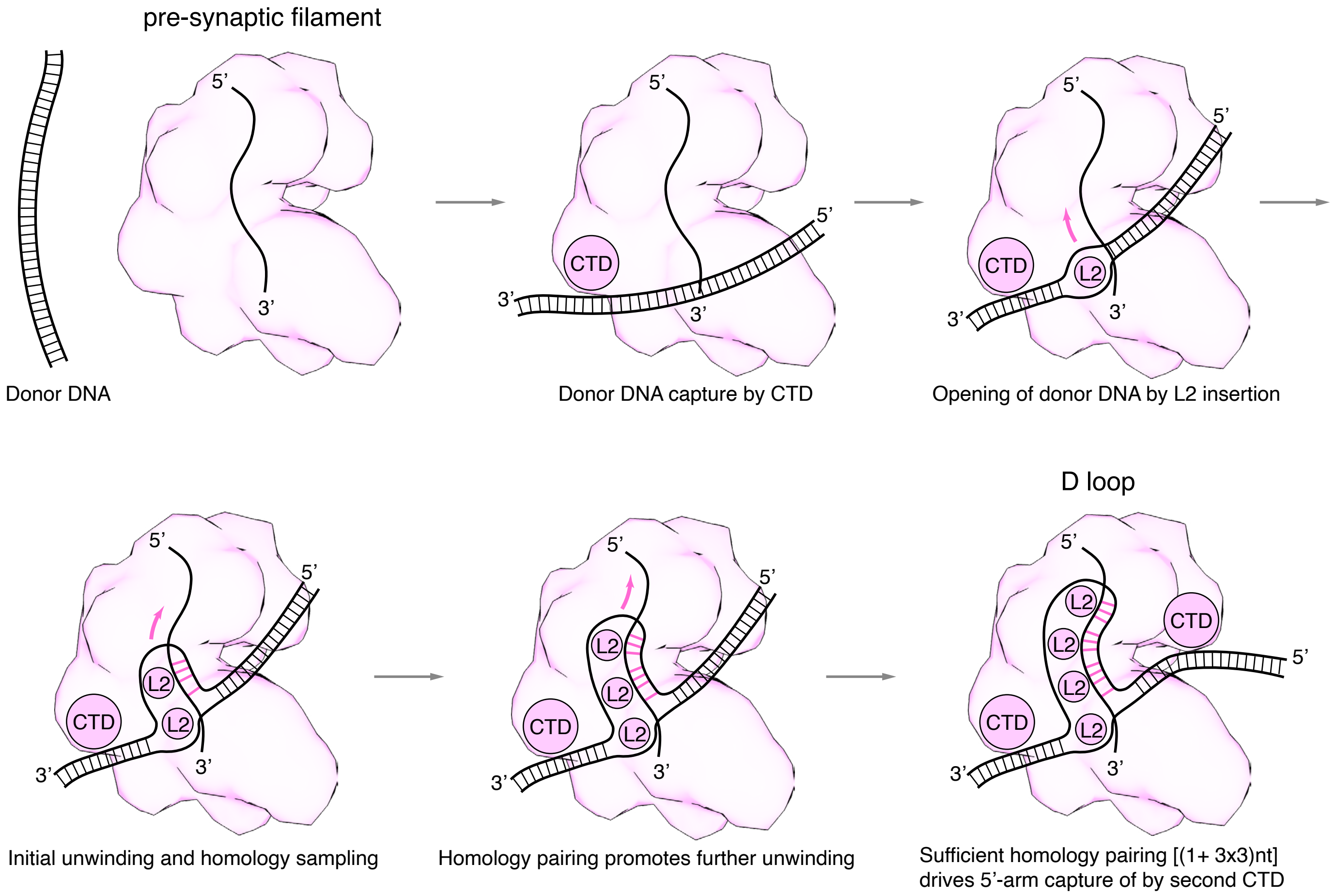


**Figure S8.** Mechanism of D-loop formation by the RecA nucleoprotein filament.

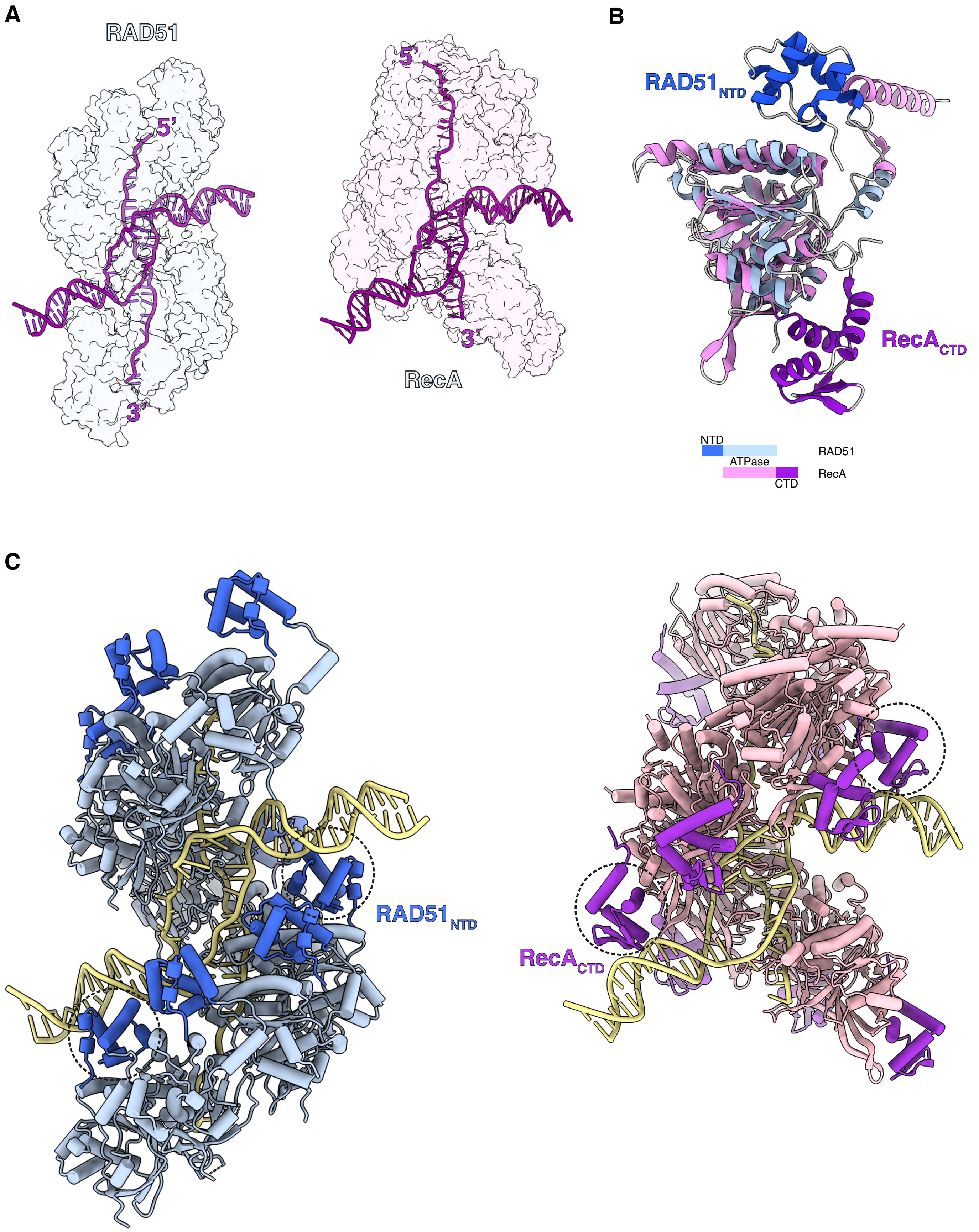


**Figure S9. Comparison of RAD51 and RecA D-loop structures. A** Side-by-side transparent surface representation of RAD51 (left, in light blue) and RecA (right, in light pink) D-loops, with DNA strands drawn as purple tubes. **B** Superposition of RAD51 and RecA protomers, coloured according to the colour key. **C** Side-by-side cylinder representation of RAD51 (left) and RecA (right) D-loops. The DNA-binding RAD51-NTD and RecA-CTD are coloured blue and magenta, respectively. The RAD51-NTD and RecA-CTD that contact the DNA are highlighted with dashed circles. The PDB accession code for the RecA D-loop is 7JY9.

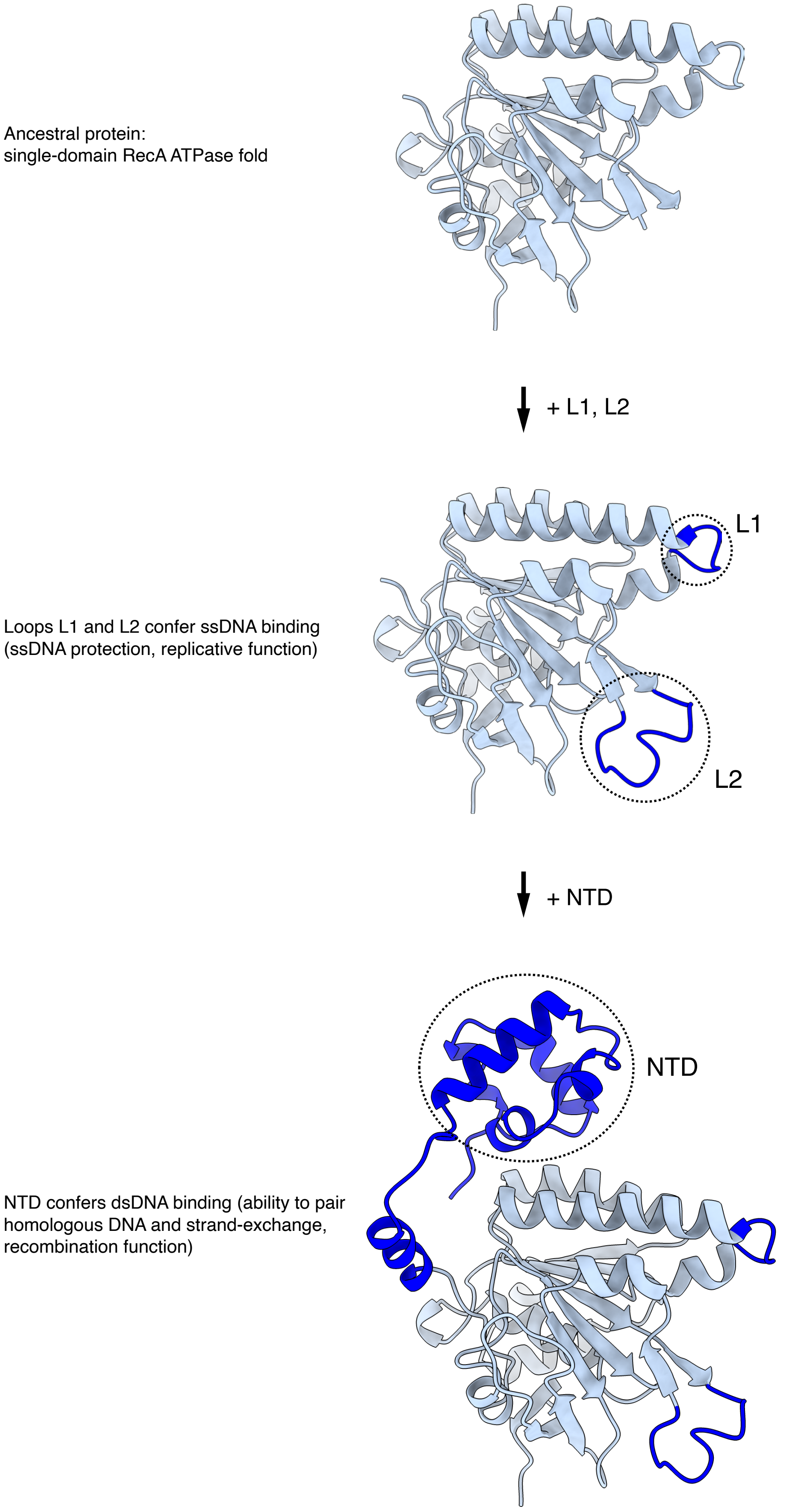


**Figure S10.** Proposed steps in the functional evolution of the RecA / RAD51 recombinases.

**Table S1.** Oligonucleotide sequence, size and labelling.

| | Size (nt) | Sequence (5'–3') |
|---|---|---|
| *EMSA & Strand Exchange* | | |
| Fig. 5B & Fig. S6B | 60 | ATGGTGTGTGTAGGTTAATGTGAGGAGGAGAGGTGAAGAAGGAGGAGAGAAGAAGGAGGC |
| Fig. 5B | 60 | ATGGTGTGTGTAGGTTAATGTGAGGAGGAGAGGTGAAGAAGGAGGAGAGAAGAAGGAGGC-**FQ** |
| Fig. 5B & Fig. S6A, B | 60 | **FAM**-GCCTCCTTCTTCTCTCCTCCTTCTTCACCTCTCCTCCTCACATTAACCTACACACACCAT |
| Fig. 5B | 60 | TTTTTTTTTTTTTTTTTTTTTTTTTTTTTTTTTTTTTTTTTTTTTTTTTTTTTTTTTTTT |
| Fig. 5D | 49 | **Alexa488**-TCAGGCGTCATTTTTCTGGTACGGAAAGTGATGCGAAAAAAACAGCGGC |
| *D-loop reconstitution and cryoEM* | | |
| Fig. S1B | 50 | TGGAGGTGCATCGAGCTCGCGACAAACCTTCTATGTTGAGCGTCAGTCGG |
| Fig. S1C | 50 | **Biotin**-TGGAGGTGCATCGAGCTCGCGACAAACCTTCTATGTTGAGCGTCAGTCGG |
| Fig. S1B | 50 | CCGACTGACGCTCAACATAGGTACCACACGGCGAGCTCGATGCACCTCCA-**Cy3** |
| Fig. S1C | 50 | **Biotin**-CCGACTGACGCTCAACATAGGTACCACACGGCGAGCTCGATGCACCTCCA-**Cy3** |
| Fig. S1B | 32 | **Cy5**-TTTTTTTTTTTCGTGTGGTACTTTTTTTTTTT |
| Fig. S1C | 32 | **Biotin**-TTTTTTTTTTTCGTGTGGTACTTTTTTTTTTT-**Biotin** |

FAM: 6-carboxyfluorescein; FQ: Iowa Black® (IDT); Cy3: cyanine 3; Cy5: cyanine 5.

### Table S2. CryoEM data collection and real-space refinement

| *Data Collection* | **Dataset #1 (free DNA)** | **Datasest #2 (streptavidin-capped DNA)** |
|---|---|---|
| Microscope | Titan Krios G3 | Titan Krios G3 |
| Voltage (keV) | 300 | 300 |
| Detector | K3 | K3 |
| Collection mode | Counting | Counting |
| Magnification | 130,000x | 130,000x |
| Pixel size (Å) | 0.326 (super-resolution) | 0.326 (super-resolution) |
| Pixel area ($Å^2$) | 0.425 | 0.425 |
| Exposure (s) | 1.31 | 1.34 |
| Dose ($e^-$/pixel/s) | 15.189 | 15.26 |
| Dose ($e^-/Å^2$/s) | 35.74 | 35.90 |
| Total dose ($e^-/Å^2$) | 46.80 | 48.10 |
| Number of frames/movie | 48 | 50 |
| Dose per frame ($e^-/Å^2$) | 0.975 | 0.96 |
| Num. movies | 2792 | 8960 |
| Defocus range (mm) | -2.6 to -1.0 (0.2) | -2.5 to -0.9 (0.2) |

*Real-space refinement*

| Composition: | |
|---|---|
| Chains | 21 |
| Non-H atoms | 24,205 |
| Residues: | |
| Protein | 2831 |
| Nucleotide | 108 |
| Ligands: | |
| ATP | 9 |
| $Ca^{2+}$ | 18 |

| Stereochemistry: | |
|---|---|
| Bonds (rmsd): | |
| Length(Å) | 0.006 |
| Angles (°) | 0.479 |
| MolProbity score[2] | 1.27 |
| Clash score | 5.04 |
| Ramachandran (%): | |
| Outliers | 0.00 |
| Allowed | 1.61 |
| Favoured | 98.39 |
| Rotamer outliers (%) | 0.49 |
| <ADP (B-factors)>: | |
| Protein | 127.59 |
| Nucleotide | 210.73 |
| Ligand | 110.90 |

| Model vs data[1]: | Resolution (Å) masked | unmasked |
|---|---|---|
| FSC, 0.143 | 2.62 | 2.65 |
| FSC, 0.5 | 2.85 | 3.06 |
| CC, mask | 0.89 | |
| CC, peaks | 0.74 | |
| CC, volume | 0.89 | |


1. Afonine PV, Klaholz BP, Moriarty NW, Poon BK, Sobolev OV, Terwilliger TC, Adams PD, Urzhumtsev A. New tools for the analysis and validation of cryo-EM maps and atomic models. Acta Crystallographica Section D: Structural Biology. 2018;74(9):814–840.

2. Chen VB, Arendall WB, Headd JJ, Keedy DA, Immormino RM, Kapral GJ, Murray LW, Richardson JS, Richardson DC. MolProbity: all-atom structure validation for macromolecular crystallography. Acta Crystallographica Section D: Biological Crystallography. 2010;66(Pt 1):12–21.
